# Common risk variants identified in autism spectrum disorder

**DOI:** 10.1101/224774

**Authors:** Jakob Grove, Stephan Ripke, Thomas D. Als, Manuel Mattheisen, Raymond Walters, Hyejung Won, Jonatan Pallesen, Esben Agerbo, Ole A. Andreassen, Richard Anney, Rich Belliveau, Francesco Bettella, Joseph D. Buxbaum, Jonas Bybjerg-Grauholm, Marie Bækved-Hansen, Felecia Cerrato, Kimberly Chambert, Jane H. Christensen, Claire Churchhouse, Karin Dellenvall, Ditte Demontis, Silvia De Rubeis, Bernie Devlin, Srdjan Djurovic, Ashle Dumont, Jacqueline Goldstein, Christine S. Hansen, Mads Engel Hauberg, Mads V. Hollegaard, Sigrun Hope, Daniel P. Howrigan, Hailiang Huang, Christina Hultman, Lambertus Klei, Julian Maller, Joanna Martin, Alicia R. Martin, Jennifer Moran, Mette Nyegaard, Terje Nærland, Duncan S. Palmer, Aarno Palotie, Carsten B. Pedersen, Marianne G. Pedersen, Timothy Poterba, Jesper B. Poulsen, Beate St Pourcain, Per Qvist, Karola Rehnström, Avi Reichenberg, Jennifer Reichert, Elise B. Robinson, Kathryn Roeder, Panos Roussos, Evald Saemundsen, Sven Sandin, F. Kyle Satterstrom, George D. Smith, Hreinn Stefansson, Kari Stefansson, Stacy Steinberg, Christine Stevens, Patrick F. Sullivan, Patrick Turley, G. Bragi Walters, Xinyi Xu, Autism Spectrum Disorders Working Group of The Psychiatric Genomics Consortium, BUPGEN, Major Depressive Disorder Working Group of the Psychiatric Genomics Consortium, 23andMe Research Team, Daniel Geschwind, Merete Nordentoft, David M. Hougaard, Thomas Werge, Ole Mors, Preben Bo Mortensen, Benjamin M. Neale, Mark J. Daly, Anders D. Børglum

**Affiliations:** The Lundbeck Foundation Initiative for Integrative Psychiatric Research, iPSYCH, Denmark; Centre for Integrative Sequencing, iSEQ, Aarhus University, Aarhus, Denmark; Department of Biomedicine - Human Genetics, Aarhus University, Aarhus, Denmark; Bioinformatics Research Centre, Aarhus University, Aarhus, Denmark; Analytic and Translational Genetics Unit, Department of Medicine, Massachusetts General Hospital and Harvard Medical School, Boston, Massachusetts, USA; Stanley Center for Psychiatric Research, Broad Institute of Harvard and MIT, Cambridge, Massachusetts, USA; Program in Medical and Population Genetics, Broad Institute of Harvard and MIT, Cambridge, Massachusetts, USA; Department of Psychiatry, Charite Universitatsmedizin Berlin Campus Benjamin Franklin, Berlin, Germany; Program in Neurogenetics, Department of Neurology, David Geffen School of Medicine, University of California, Los Angeles, USA; Center for Autism Research and Treatment and Center for Neurobehavioral Genetics, Semel Institute for Neuroscience and Human Behavior, University of California, Los Angeles, USA; National Centre for Register-Based Research, Aarhus University, Aarhus, Denmark; Centre for Integrated Register-based Research, Aarhus University, Aarhus Denmark; NORMENT - KG Jebsen Centre for Psychosis Research, University of Oslo, Oslo, Norway; Division of Mental Health and Addiction, Oslo University Hospital, Oslo, Norway; MRC Centre for Neuropsychiatric Genetics and Genomics, Cardiff University, Cardiff, UK; Seaver Autism Center for Research and Treatment, Icahn School of Medicine at Mount Sinai, New York, USA; Department of Psychiatry, Icahn School of Medicine at Mount Sinai, New York, NY 10029, USA; Friedman Brain Institute, Icahn School of Medicine at Mount Sinai, New York, NY 10029, USA.; Center for Neonatal Screening, Department for Congenital Disorders, Statens Serum Institut, Copenhagen, Denmark; Department of Medical Epidemiology and Biostatistics, Karolinska, Sweden; Department of Psychiatry, University of Pittsburgh School of Medicine, Pittsburgh, PA 15213, USA.; Department of Medical Genetics, Oslo University Hospital, Oslo, Norway; Institute of Biological Psychiatry, MHC Sct. Hans, Mental Health Services Copenhagen, Denmark; Department of Neurohabilitation, Oslo University Hospital, Oslo; Norway; Genomics plc, Oxford, UK; Department of Rare Disorders and Disabilities, Oslo University Hospital, Norway; Institute for Molecular Medicine Finland, University of Helsinki, Helsinki, Finland.; Language and Genetics Department, Max Planck Institute for Psycholinguistics, Nijmegen, The Netherlands; MRC Integrative Epidemiology Unit, University of Bristol, Bristol, UK; Donders Institute for Brain, Cognition and Behaviour, Radboud University, Nijmegen, The Netherlands; Wellcome Trust Sanger Institute, Hinxton, Cambridge CB10 1SA, UK; Department of Epidemiology, Harvard Chan School of Public Health, Boston, Massachusetts, USA; Computational Biology Department, Carnegie Mellon University, Pittsburgh, PA 15213, USA; Department of Statistics, Carnegie Mellon University, Pittsburgh, PA 15213, USA.; Institute for Genomics and Multiscale Biology, Department of Genetics and Genomic Sciences, Icahn School of Medicine at Mount Sinai, New York, NY, USA; Mental Illness Research Education and Clinical Center (MIRECC), James J. Peters VA Medical Center, Bronx, New York, USA; The State Diagnostic and Counselling Centre, Digranesvegur 5, IS-200 Kópavogur, Iceland; School of Social and Community Medicine, University of Bristol, Bristol, UK; deCODE genetics/Amgen, Sturlugata 8, IS-101 Reykjavík Iceland; Departments of Genetics and Psychiatry, University of North Carolina, Chapel Hill, NC, USA; Faculty of Medicine, University of Iceland, Reykjavik, Iceland; Department of Human Genetics, David Geffen School of Medicine, University of California, Los Angeles, California, USA; Mental Health Services in the Capital Region of Denmark, Mental Health Center Copenhagen, University of Copenhagen, Copenhagen, Denmark; Department of Clinical Medicine, University of Copenhagen, Copenhagen, Denmark; Psychosis Research Unit, Aarhus University Hospital, Risskov, Denmark

## Abstract

Autism spectrum disorder (ASD) is a highly heritable and heterogeneous group of neurodevelopmental phenotypes diagnosed in more than 1% of children. Common genetic variants contribute substantially to ASD susceptibility, but to date no individual variants have been robustly associated with ASD. With a marked sample size increase from a unique Danish population resource, we report a genome-wide association meta-analysis of 18,381 ASD cases and 27,969 controls that identifies five genome-wide significant loci. Leveraging GWAS results from three phenotypes with significantly overlapping genetic architectures (schizophrenia, major depression, and educational attainment), seven additional loci shared with other traits are identified at equally strict significance levels. Dissecting the polygenic architecture we find both quantitative and qualitative polygenic heterogeneity across ASD subtypes, in contrast to what is typically seen in other complex disorders. These results highlight biological insights, particularly relating to neuronal function and corticogenesis and establish that GWAS performed at scale will be much more productive in the near term in ASD, just as it has been in a broad range of important psychiatric and diverse medical phenotypes.

---

ASD is the term for a group of pervasive neurodevelopmental disorders characterized by impaired social and communication skills along with repetitive and restrictive behavior. The clinical presentation is very heterogeneous including individuals with severe impairment and intellectual disability as well as individuals with above average IQ and high levels of academic and occupational functioning. ASD affects 1-1.5% of individuals and is highly heritable, with both common and rare variants contributing to its etiology^1–4^. Common variants have been estimated to account for a major part of ASD liability^2^ as has been observed for other common neuropsychiatric disorders. By contrast, *de novo* mutations, mostly copy number variants (CNVs) and gene disrupting point mutations, have larger individual effects, but collectively explain <5% of the overall liability^1–3^ and far less of the heritability. While a number of genes have been convincingly implicated via excess statistical aggregation of *de novo* mutations, the largest GWAS to date (n=7387 cases scanned) – while providing compelling evidence for the bulk contribution of common variants – did not conclusively identify single variants at genome-wide significance^5–7^. This underscored that common variants, as in other complex diseases such as schizophrenia, individually have low impact and that a substantial scale-up in sample numbers would be needed.

Here we report the first common risk variants robustly associated with ASD by more than doubling the discovery sample size compared to previous GWAS^5–8^. We describe strong genetic correlations between ASDs and other complex disorders and traits, confirming shared etiology, and we show results indicating differences in the polygenic architecture across clinical sub-types of ASD. Leveraging these relationships and recently introduced computational techniques^9^, we identify additional novel ASD-associated variants that are shared with other phenotypes. Furthermore, by integrating with complementary data from Hi-C chromatin interaction analysis of fetal brains and brain transcriptome data, we explore the functional implications of our top-ranking GWAS results.

## Results

### GWAS

As part of the iPSYCH project (http://ipsych.au.dk)^10^, we collected and genotyped a Danish nation-wide population-based case-cohort sample including nearly all individuals born in Denmark between 1981 and 2005 and diagnosed with ASD (according to ICD-10) before 2014. We randomly selected controls from the same birth cohorts (**Table S1.1.1**). We have previously validated registry-based ASD diagnoses^11,12^ and demonstrated the accuracy of genotyping DNA extracted and amplified from bloodspots collected shortly after birth^13,14^. Genotypes were processed using Ricopili^15^, performing stringent quality control of data, removal of related individuals, exclusion of ancestry outliers based on principal component analysis, and imputation^16,17^ using the 1000 Genomes Project phase 3 reference panel. After this processing, genotypes from 13,076 cases and 22,664 controls from the iPSYCH sample were included in analysis. As is now standard in human complex trait genomics, our primary analysis is a meta-analysis of the iPSYCH ASD results with five family-based trio samples of European ancestry from the Psychiatric Genomics Consortium (PGC, 5,305 cases and 5,305 pseudo controls^18^. All PGC samples had been processed using the same Ricopili pipeline for QC, imputation and analysis as employed here.

Supporting the consistency between the study designs, the iPSYCH population-based and PGC family-based analyses showed a high degree of genetic correlation with *r_G_* = 0.779 (SE =0.106; P = 1.75 x 10^−13^), similar to the genetic correlations observed between datasets in other mental disorders^19^. Likewise, polygenicity as assessed by polygenic risk scores (PRS)^20^ showed consistency across the samples supporting homogeneity of effects across samples and study designs (**Figure S4.4.4**). SNP heritability 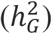 was estimated to be 0.118 (SE = 0.010, for population prevalence of 0.012 ^21^).

The main GWAS meta-analysis totaled 18,381 ASD cases and 27,969 controls, and applied an inverse variance-weighted fixed effects model^22^. To ensure that the analysis was well-powered and robust, we examined markers with minor allele frequency (MAF) ≥ 0.01, imputation INFO score ≥ 0.7, and supported by an effective sample size in >70% of the total. This final meta-analysis included results for 9,112,387 autosomal markers and yielded 93 genome-wide significant markers in three separate loci (**Figure 1**; **Table 1a**; **Figures S4.2.1-4.2.44**). Each locus was strongly supported by both the Danish case-control and the PGC family-based data. While modest inflation was observed (lambda=1.12, lambda 1000 = 1.006), LD score regression analysis^23^ indicates this is arising from polygenicity (> 96%, see Methods) rather than confounding. The strongest signal among 265,846 markers analyzed on chromosome X was P = 5.4 x 10^−6^.

**Figure 1.**
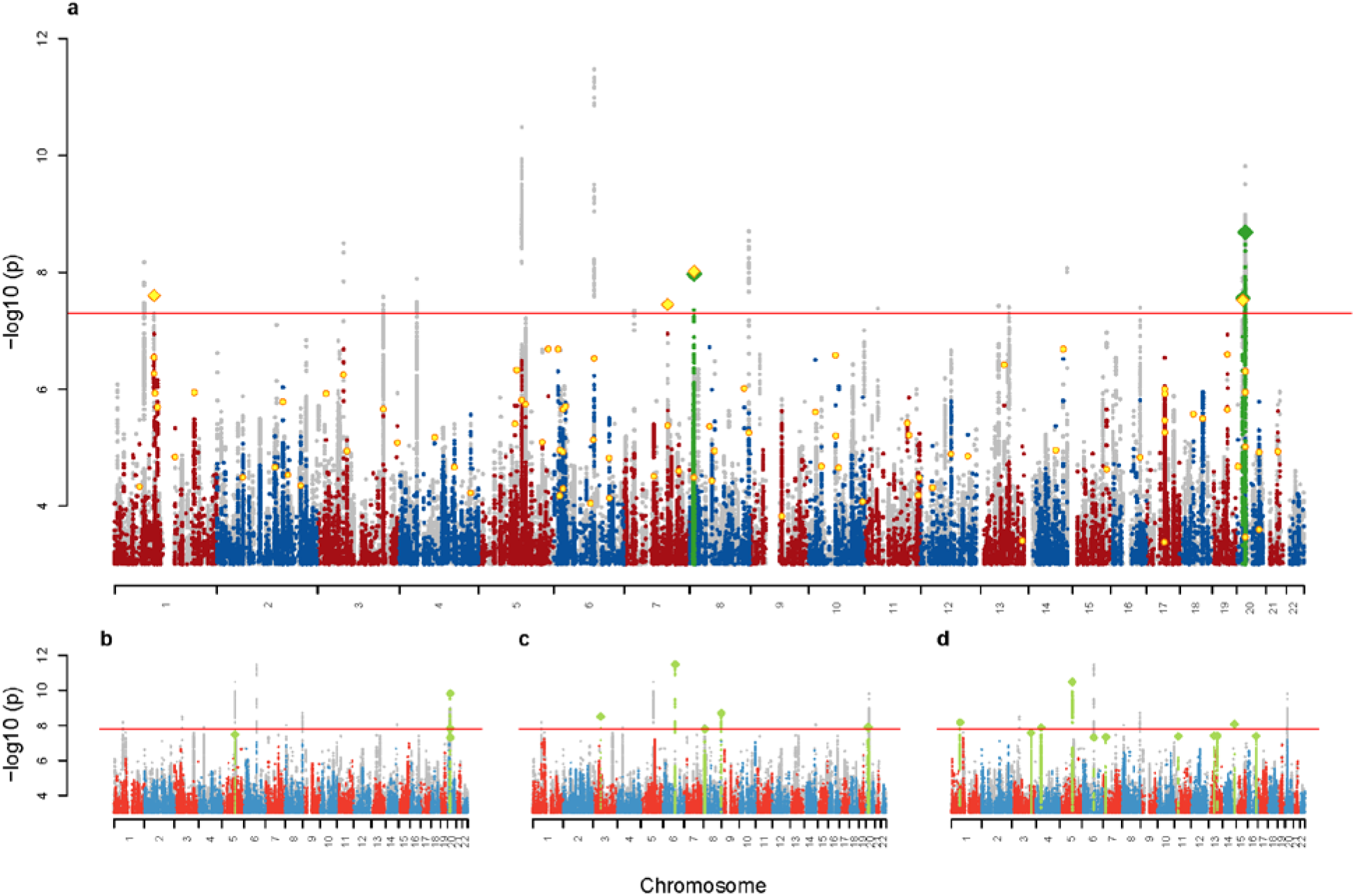
Manhattans plots. **a**: The main ASD scan with the results of the combined analysis with the follow-up sample in yellow in the foreground. GWS clumps are painted green with index SNPs as diamonds. **b-d**: Manhattan plots for three MTAG scans of ASD together with, respectively, schizophrenia^15^, educational attainment^26^ and major depression^29^ (see Figures **S.2.71-74** for full size plots). In all panels the results of the composite of the five analyses (consisting for each marker of the minimal p-value of the five) is shown in grey in the background.

**Table 1.**
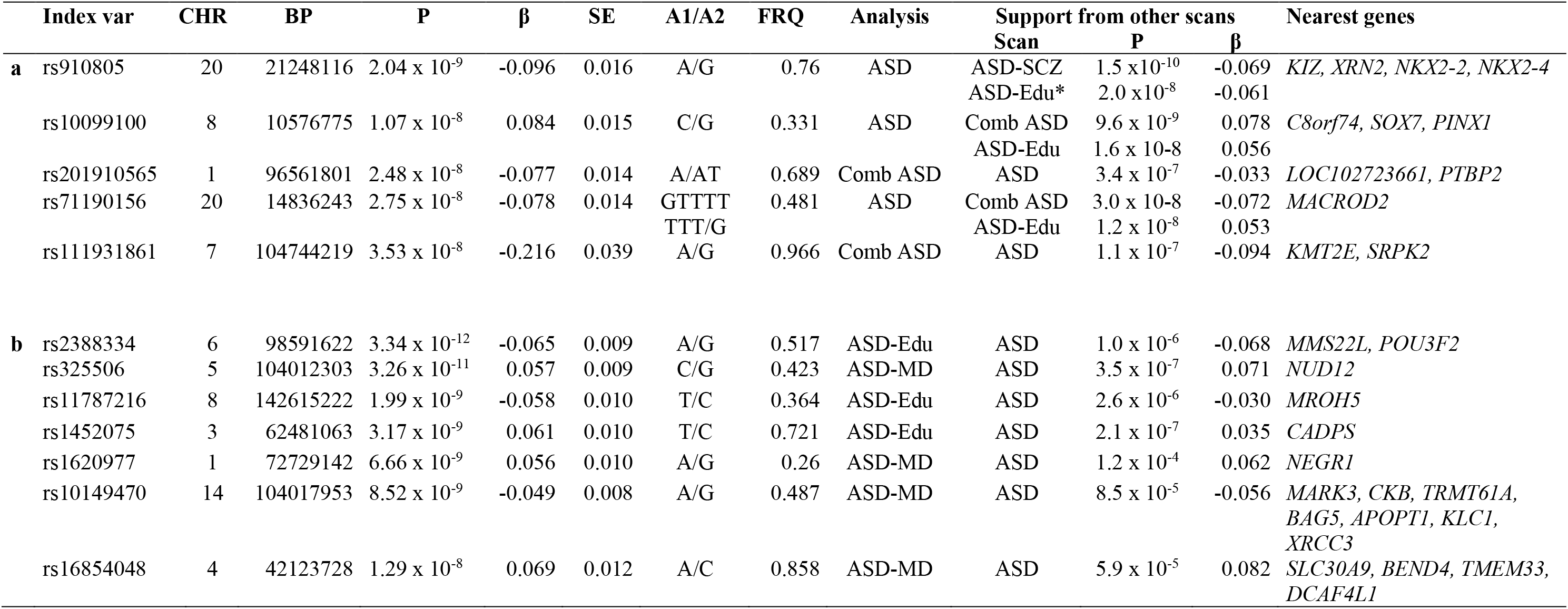
Genome-wide significant loci. Independent loci (r^2^<0.1, distance > 400kb) with index variant (Index var), chromosome (CHR), chromosomal position (BP), alleles (A1/A2), allele frequency of A1 (FRQ), estimate of effect (β) with respect to A1, standard error of β (SE), and the association p-value of the index variant (P). The “Analysis” column shows the analysis that generated the particular result. The column “Support from other scans” lists the analyses that also support the locus. For the ASD scan results, this shows the genome wide significant results in the locus from the other scans, and for results from any other analysis, it shows the result from the ASD scan. An * indicate different lead SNP. The column “Nearest genes” lists nearest genes from within 50kb of the r^2^≥0.6 LD friends of the index variant (intergenic variants marked with a dash). **a.** Loci from the ASD scan and the combined analysis with the follow-up sample. Column “Analysis” indicates which results stem from the original scan and which comes from the combined. **b.** Loci from the three MTAG analyses with schizophrenia (SCZ)^15^, educational attainment (Edu)^26^ and major depression (MD)^29^ – loci not already represented in section a.

We next obtained replication data for the top 88 loci with p-values < 1x10^−5^ in five cohorts of European ancestry totaling 2,119 additional cases and 142,379 controls (**Table S1.3.2**). An overall replication of direction of effects was observed (53 of 88 (60%) of P < 1x10^−5^; 16 of 23 (70%) at P < 1x10^−6^; both sign tests P < 0.05) and two additional loci achieved genome-wide significance in the combined analysis (**Table 1a**). More details on the identified loci can be found in **Table S3.1.3** and selected candidates are described in **Box1**.

**Box1.** Selected loci and candidates (ordered by chromosome).

| Gene | Locus* and supporting evidence | Gene function |
| --- | --- | --- |
| NEGR1 | Chr1:72,729,142<br>Shared ASD-MDD locus<br>Locus also significant in depression<sup>29,78</sup>, obesity and BMI<sup>84-88</sup><br>NEGR1 is the only protein-coding gene in the locus<br>NEGR1 is supported by brain Hi-C and eQTL analyses<sup>29</sup> | NEGR1 (neuronal growth regulator 1) is an adhesion molecule modulating synapse formation in hippocampal neurons<sup>89,90</sup> and neurite outgrowth<sup>91,92</sup>. It is member of the IgLON protein family implicated in synaptic plasticity and axon extension<sup>93-95</sup>.<br>Predominantly expressed (and developmentally upregulated) in hippocampus and cortex<sup>96</sup> and also hypothalamus<sup>97</sup>. |
| PTBP2 | Chr1:96,561,801<br>ASD locus<br>Locus also significant in BMI<sup>84,86,88</sup> and weight<sup>84</sup>. In schizophrenia the locus show p-value of $6.5 \times 10^{-6}$<sup>15</sup><br>PTBP2 is the nearest protein-coding gene, approx. 625kb from index SNP.<br>De novo and rare variants in PTBP2 have been reported in ASD cases<sup>1,3,98</sup>.<br>PTBP2 is supported by Hi-C results in this study (Fig. 5d) | PTBP2 is also known as nPTB (neuronal PTB) or brPTB (brain PTB) and is a splicing regulator. PTBP1 and its paralog PTBP2 bind to intronic polypyrimidine tracts in pre-mRNAs and target large sets of exons to coordinate alternative splicing programs during development<sup>99</sup>. Several switches in the expression of PTBP1 and PTBP2 regulate alternative splicing during neurogenesis and neuronal differentiation<sup>100-103</sup>. |
| CADPS | Chr3:62,481,063<br>Shared ASD-Educational attainment locus<br>Locus also significant in study of cognitive decline rate<sup>104</sup><br>CADPS is supported by Hi-C results in this study (Fig. 5a). | CADPS encodes a calcium-binding protein involved in exocytosis of neurotransmitters and neuropeptides. In line with CAPDS mRNA being mainly expressed in brain and pituitary (gtexportal), immunoreactive CAPS-1 is localized in neural and various endocrine tissues<sup>105</sup>. In hippocampal synapses, CADPS regulates the pool of readily releasable vesicles at pre-synaptic terminals<sup>106,107</sup> |
| KCNN2 | Chr5: 113,801,423<br>ASD locus (gene-wise analysis)<br>Locus also significant in educational attainment<sup>26,108</sup>.<br>KCNN2 synaptic levels are regulated by the E3 ubiquitin ligase UBE3A<sup>109</sup>, of which overexpression has been linked to ASD risk<sup>109,110</sup>. | KCNN2 is a voltage-independent Ca<sup>2+</sup>-activated K<sup>+</sup> channel that responds to changes in intracellular calcium concentration and couples calcium metabolism to potassium flux and membrane excitability. In CNS neurons, activation of KCNN2 modulates neuronal excitability by causing membrane hyperpolarization<sup>111</sup>. Hippocampal KCNN2 has roles in the formation of new memory<sup>112</sup>, encoding and consolidation of contextual fear<sup>113</sup>, and in drug-induced plasticity<sup>114</sup>. |
| KMT2E | Chr7:104,744,219<br>ASD locus<br>Locus also significant in schizophrenia<sup>15,115</sup> and in ASD-schizophrenia meta-analysis<sup>5</sup>.<br>KMT2E de novo mutations are associated with ASD at FDR<0.1<sup>116</sup><br>A KMT2E credible SNP is a loss-of-function variant (Table S3.4.1) | KMT2E encodes Histone-lysine N-methyltransferase 2E and forms a family together with SETD5<sup>117,118</sup>. Evidence suggest that recognition of the histone H3K4me3 mark by the KMT2E PHD finger can facilitate the recruitment of KMT2E to transcription-active chromatin regions<sup>119,120</sup>.<br>KMT2E has been implicated in chromatin regulation, control of cell cycle progression, and maintaining genomic stability<sup>121</sup>. |
| MACROD2 | Chr20: 14836243<br>ASD locus<br>Locus found significant in previous ASD GWAS<sup>61</sup> but not supported in larger study<sup>122</sup><br>MACROD2 is the only protein-coding gene in the locus | MACROD2 is a nuclear enzyme that binds to mono-ADP-ribosylated (MARylated) proteins and functions as an eraser of mono-ADP-ribosylation<sup>123</sup>. Intracellular MARylated histones and GSK3β are substrates of MACROD2, and the removal of MAR from GSK3β is responsible for reactivating of its kinase activity<sup>123</sup>. This gene is expressed in lung and multiple regions of the brain. Low or no expression across most other tissue (https://www.gtexportal.org). |

\*position of index SNP is listed.

### Correlation with other traits and multi-trait GWAS

To investigate the extent of genetic overlap between ASD and other phenotypes we estimated the genetic correlations with a broad set of psychiatric and other medical diseases, disorders, and traits available at LD Hub^24,25^ using bivariate LD score regression (**Figure 2**, **Table S3.3.2**). Significant correlations were found for several traits including schizophrenia^15^ (*r_G_* = 0.211, p = 1.03 x 10^−5^) and measures of cognitive ability, especially educational attainment^26^ (*r_G_* = 0.199, p = 2.56 x 10^−9^), indicating a substantial genetic overlap with these phenotypes and corroborating previous reports^5,24,27,28^. In contrast to previous reports^18^, we found a strong and highly significant correlation with major depression^29^ (*r_G_* = 0.412, p = 1.40 x 10^−25^), and we are the first to report a prominent overlap with ADHD^30^ (*r_G_* = 0.360, p = 1.24 x 10^−12^). Moreover, we confirm the genetic correlation with social communication difficulties at age 8 in a non-ASD population sample reported previously based on a subset of the ASD sample^31^ (*r_G_* = 0.375, p = 0.0028).

**Figure 2.**
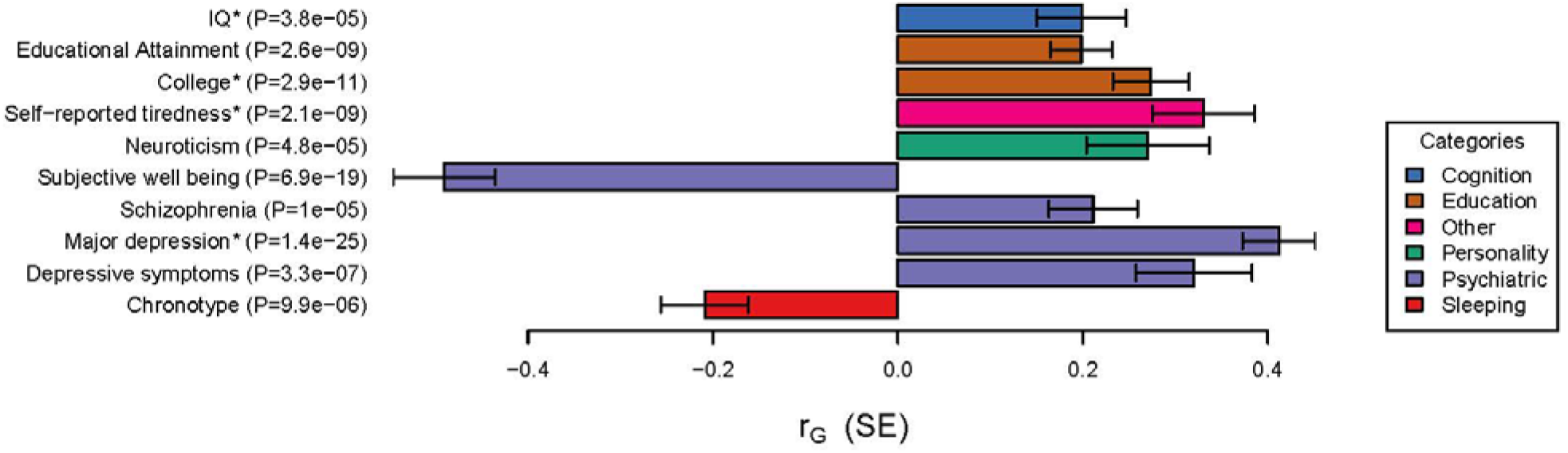
Genetic correlation with other traits. Significant genetic correlations between ASD and other traits after Bonferroni correction for testing a total of 234 traits available at LDhub with the addition of a handful of new phenotypes. The results here corresponds to the following GWAS analyses: IQ^43^, educational attainment^26^, college^124^, self-reported tiredness^125^, neuroticism^27^, subjective well-being^27^, schizophrenia^15^, major depression^29^, depressive symptoms^27^, ADHD^30^, and chronotype^44^. See Table **S3.3.2** for the full output of this analysis. * Indicates that the values are from in-house analyses of new summary statistics not yet included in LD Hub.

In order to leverage these observations for the discovery of ASD loci shared with these other traits, we selected three particularly well-powered and genetically correlated phenotypes. These were schizophrenia (N = 79,641)^15^, major depression (N = 424,015)^29^ and educational attainment (N = 328,917)^26^. We utilize the recently introduced MTAG method^9^ which, briefly, generalizes the standard inverse-variance weighted meta-analysis for multiple phenotypes. In this case, MTAG takes advantage of the fact that, given an overall genetic correlation between ASD and a second trait, the effect size estimate and evidence for association to ASD can be improved by appropriate use of the association information from the second trait. The results of these three ASD-anchored MTAG scans are correlated to the primary ASD scan (and to each other) but given the exploration of three scans, we utilize a more conservative threshold of 1.67 x 10^−8^ for declaring significance across these secondary scans. In addition to stronger evidence for several of the ASD hits defined above, variants in seven additional regions achieve genome-wide significance, including three loci shared with educational attainment and four shared with major depression (**Table 1b, Box 1**).

### Gene and gene-set analysis

Next, we performed gene-based association analysis on our primary ASD meta-analysis using MAGMA^32^, testing for the joint association of all markers within a locus (across all protein-coding genes in the genome). This analysis identified 15 genes surpassing the significance threshold (**Table S3.1.2**). As expected, the majority of these genes were located within the genome-wide significant loci identified in the GWAS, but seven genes are located in four additional loci including *KCNN2*, *MMP12*, *NTM* and a cluster of genes on chromosome 17 (*KANSLl*, *WNT3*, *MAPT* and *CRHRl*) (**Figures S4.2.46-60**). In particular, *KCNN2* was strongly associated (P = 1.02x10^−9^), far beyond even single-variant statistical thresholds and is included in the descriptions in **Box 1**.

Enrichment analyses using gene co-expression modules from human neocortex transcriptomic data (M13, M16 and M17 from Parikshak et al. 2013 ^33^ and loss-of-function intolerant genes (pLI > 0.9)^34,35^, which previously have shown evidence of enrichment in neurodevelopmental disorders^30,33,36^, yielded only nominal significance for the latter and M16 (**Table S3.1.4**). Likewise analysis of Gene Ontology sets^37,38^ for molecular function from MsigDB^39^ revealed no significant sets after Bonferroni correction for multiple testing (**Table S3.1.5**).

### Dissection of the polygenic architecture

As ASD is a highly heterogeneous disorder, we explored how 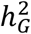 partitioned across phenotypic sub-categories in the iPSYCH sample and estimated the genetic correlations between these groups using GCTA^40–42^. We examined cases with and without intellectual disability (ID, N = 1,873) and the ICD-10 diagnostic sub-categories: childhood autism (F84.0, N = 3,310), atypical autism (F84.1, N = 1,607), Asperger’s syndrome (F84.5, N = 4,622), and other/unspecified pervasive developmental disorders (PDD, F84.8-9, N = 5,795), reducing to non-overlapping groups when doing pairwise comparisons (see **Table S2.3.1**). While the pairwise genetic correlations were consistently high between all sub-groups (95% CIs including 1 in all comparisons), the 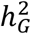 of Asperger’s syndrome (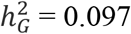, SE = 0.001) was found to be twice the 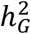 of both childhood autism (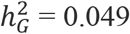, SE =0.009, P =0.001) and the group of other/unspecified PDD (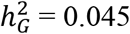, SE =0.008, P =0.001) (**Table S3.2.1** and **S3.3.1, Figure S4.3.1** and **S4.4.1**). Similarly, the 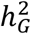 of ASD without ID (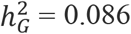, SE = 0.005) was found to be three times higher than for cases with ID (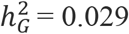, SE = 0.013, P = 0.015).

To further examine the apparent polygenic heterogeneity across subtypes, we investigated how PRS trained on different phenotypes were distributed across distinct ASD subgroups. We focused on phenotypes showing strong genetic correlation with ASD (e.g., educational attainment), but included also traits with little or no correlation to ASD (e.g., BMI) as negative controls. In this analysis, we regressed the normalized scores on ASD subgroups while including covariates for batches and principal components in a multivariate regression. Of the eight phenotypes we evaluated, only the cognitive phenotypes showed strong heterogeneity (educational attainment^26^ P = 1.8 x 10^−8^, IQ^43^ P = 3.7 x 10^−9^) (**Figure S4.4.8**). Interestingly, all case-control groups with or without ID showed significantly different loading for the two cognitive phenotypes: controls with ID have the lowest score followed by ASD cases with ID, and ASD cases without ID have significantly higher scores again than any other group (P = 2.6 x 10^−12^ for educational attainment, P = 8.2 x 10^−12^ for IQ).

With respect to the diagnostic sub-categories constructed hierarchically from ASD subtypes (**Table S2.3.1**), it was again the cognitive phenotypes that showed the strongest heterogeneity across the diagnostic classes (educational attainment P = 2.6 x 10^−11^, IQ P = 3.4 x 10^−8^), while neuroticism^27^ (P = 0.0015), chronotype^44^ (P = 0.011) and subjective wellbeing^27^(P = 0.029) showed weaker but nominally significant degree of heterogeneity, and SCZ, MDD and BMI were non-significant across the groups (P > 0.19) (**Figure 3**). This pattern weakened only slightly when excluding subjects with ID (**Figure S4.4.9**). For neuroticism there was a clear split with atypical and other/unspecified PDD cases having significantly higher PRS than childhood autism and Asperger’s, P =0.00013. Considering the genetic overlap of each subcategory with each phenotype, the hypothesis of homogeneity across sub-phenotypes was strongly rejected (P = 1.6 x 10^−11^), thereby establishing that these sub-categories indeed have differences in their genetic architectures.

**Figure 3.**
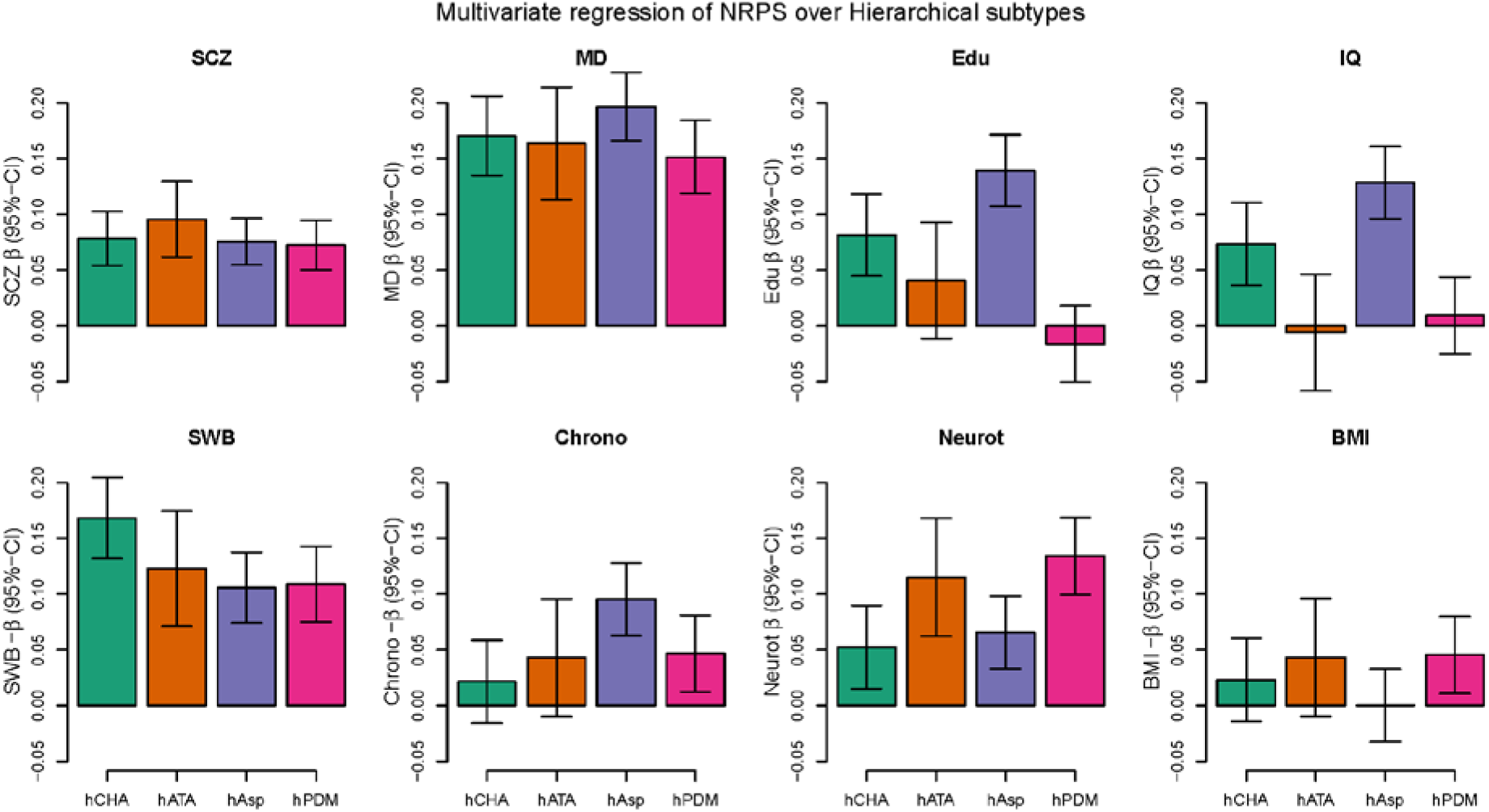
Profiling PRS load across distinct ASD sub-groups for 8 different phenotypes. (schizophrenia^15^, major depression^29^, educational attainment^26^, human intelligence^43^, subjective well-being^27^, chronotype^44^, neuroticism^27^ and BMI^88^. The bars show coefficients from multivariate regression of the 8 normalized scores on the distinct ASD sub-types (adjusting for batches and PCs). The subtypes are the hierarchically defined subtypes for childhood autism (hCHA), atypical autism (hATA), Asperger’s (hAsp), and the lumped pervasive disorders developmental group (hPDM). Beware that the orientation of the scores for subjective well-being, chronotype and BMI have been switched to improve graphical presentation. The corresponding plot where subjects with ID have been excluded can be seen in Figure **S4.4.9**, and with ID as a subtype in Figure **S4.4.8**. Applying the same procedure to the internally trained ASD score did not display systematic heterogeneity (p=0.068) except as expected for the ID groups (p=0.00027) (**Figure S4.4.10**).

Focusing on educational attainment, significant enrichment of PRS was found for Asperger’s syndrome (P = 2.0 x 10^−17^) in particular, and for childhood autism (P = 1.5 x 10^−5^), but not for the group of other/unspecified PDD (P =0.36) or for atypical autism (P =0.13) (**Figure 3**). Excluding individuals with ID only changes this marginally, with atypical autism becoming nominally significant (P =0.020) (**Figure S4.4.9**). These results reveal that the genetic architecture underlying educational attainment is indeed shared with ASD but to a variable degree across the disorder spectrum. We find that the observed excess in ASD subjects of alleles positively associated with education attainment^45,46^ is confined to Asperger’s and childhood autism, and it is not seen here in atypical autism nor in other/unspecified PDD.

Finally, we evaluated the predictive ability of ASD PRS using five different sets of target and training samples within the combined iPSYCH-PGC sample. The observed mean variance explained by PRS (Nagelkerke’s R^2^) was 2.45% (P = 5.58 x 10^−140^) with a pooled PRS-based case-control odds ratio OR = 1.33 (CI.95% 1.30 – 1.36) (**Figures S4.4.2** and **S4.4.4**). Dividing the target samples into PRS decile groups revealed an increase in OR with increasing PRS. The OR for subjects with the highest PRS increased to OR = 2.80 (CI.95%2.53–3.10) relative to the lowest decile (**Figures 4a and S4.4.5**). Leveraging correlated phenotypes in an attempt to improve prediction of ASD, we generated a multi-phenotype PRS as a weighted sum of phenotype specific PRS (see Methods). As expected, Nagelkerkes’s R^2^ increased for each PRS included attaining its maximum at the full model at 3.77% (P = 2.03 x 10^−215^) for the pooled analysis with an OR = 3.57 (CI.95% 3.22–3.96) for the highest decile (**Figures 4b and S4.4.6-7**). These results demonstrate that an individual’s ASD risk depends on the level of polygenic burden of thousands of common variants in a dose-dependent way, which can be reinforced by adding SNP-weights from ASD correlated traits.

**Figure 4.**
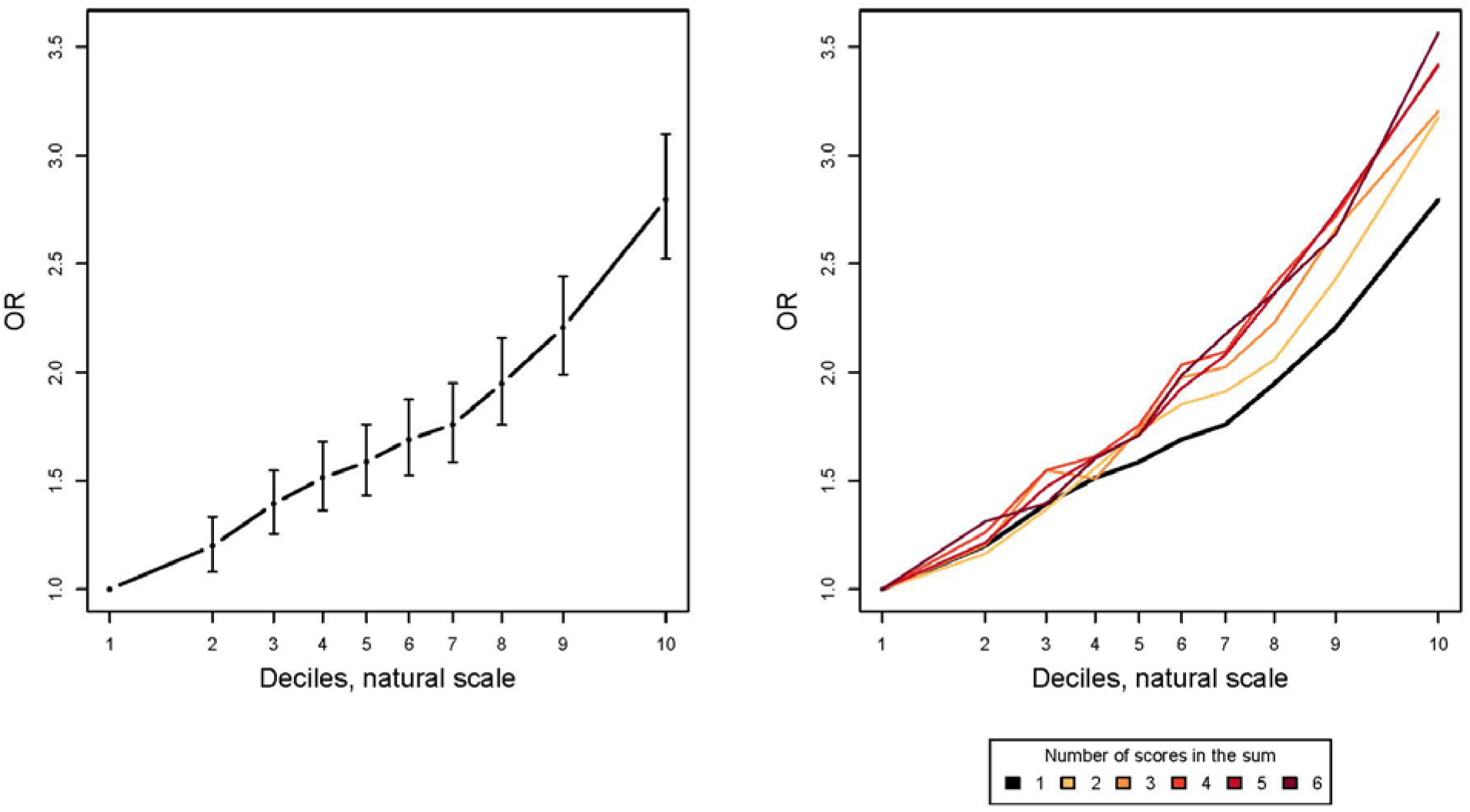
Decile plots. (Odds Ratio (OR) by PRS within each decile): **a.** Decile plot with 95%-CI for the internally trained ASD score (P-value threshold is 0.1). **b.** Decile plots on a weighted sums of PRSs starting with the ASD score of panel a and successively adding the scores for major depression^29^, subjective well-being^27^, schizophrenia^15^, educational attainment^26^, and chronotype^44=44^. In all instances the P-value threshold for the score used is the one with the highest Nagelkerke’s *R*^2^. Figures **S4.4.5** and **S4.4.7** show the stability across leave-one out groups that was used to create these combined results.

### Functional annotation

In order to obtain information on possible biological underpinnings of our GWAS results we conducted several analyses. First, we examined how the ASD h£ partitioned on functional genomic categories as well as on cell type-specific regulatory elements using stratified LD score regression^47^. This analysis revealed significant enrichment of heritability in conserved DNA regions and H3K4me1 histone marks^48^, as well as in genes expressed in central nervous system (CNS) cell types as a group (**Figures S4.3.2** and **S4.3.3**), which is in line with observations in schizophrenia, major depression^29^, and bipolar disorder^24^. Analyzing the enhancer associated mark H3K4me1 in individual cell/tissue^48^, we found significant enrichment in brain and neuronal cell lines (**Figure S4.3.4**). The highest enrichment was observed in the developing brain, germinal matrix, cortex-derived neurospheres, and embryonic stem cell (ESC)-derived neurons, consistent with ASD as a neurodevelopmental disorder with largely prenatal origins, as supported by data from analysis of rare *de novo* variants^33^.

Common variation in ASD is located in regions that are highly enriched with regulatory elements predicted to be active in human corticogenesis (**Figures S4.3.2-4**). Since most gene regulatory events occur at a distance via chromosome looping, we leveraged Hi-C data from germinal zone (GZ) and post-mitotic zones cortical plate (CP) in the developing fetal brain to identify potential target genes for these variants^49^. We performed fine mapping of 29 loci to identify the set of credible variants containing likely causal genetic risk^50^ (see Methods). Credible SNPs were significantly enriched with enhancer marks in the fetal brain (Figure S4.5.1), again confirming the likely regulatory role of these SNPs during brain development.

Based on location or evidence of physical contact from Hi-C, the 380 credible SNPs (29 loci) could be assigned to 95 genes (40 protein-coding), including 39 SNPs within promoters that were assigned to 9 genes, and 16 SNPs within the protein coding sequence of 8 genes (**Table S3.4.1, Figure S4.5.1**). Hi-C identified 86 genes, which interacted with credible SNPs in either the CP or GZ during brain development. Among these genes, 34 are interacting with credible SNPs in both CP and GZ, which represent a high-confidence gene list. Notable examples are illustrated in **Figure 5** and highlighted in **Box 1**. By analyzing their mean expression trajectory, we observed that the identified ASD candidate genes (**Table S3.4.1**) show highest expression during fetal corticogenesis, which is in line with the enrichment of heritability in the regulatory elements in developing brain (**Figure 5e-g**). Interestingly, both common and rare variation in ASD preferentially affects genes expressed during corticogenesis^33^, highlighting a potential spatiotemporal convergence of genetic risk on this specific developmental epoch, despite the disorder’s profound genetic heterogeneity.

**Figure 5.**
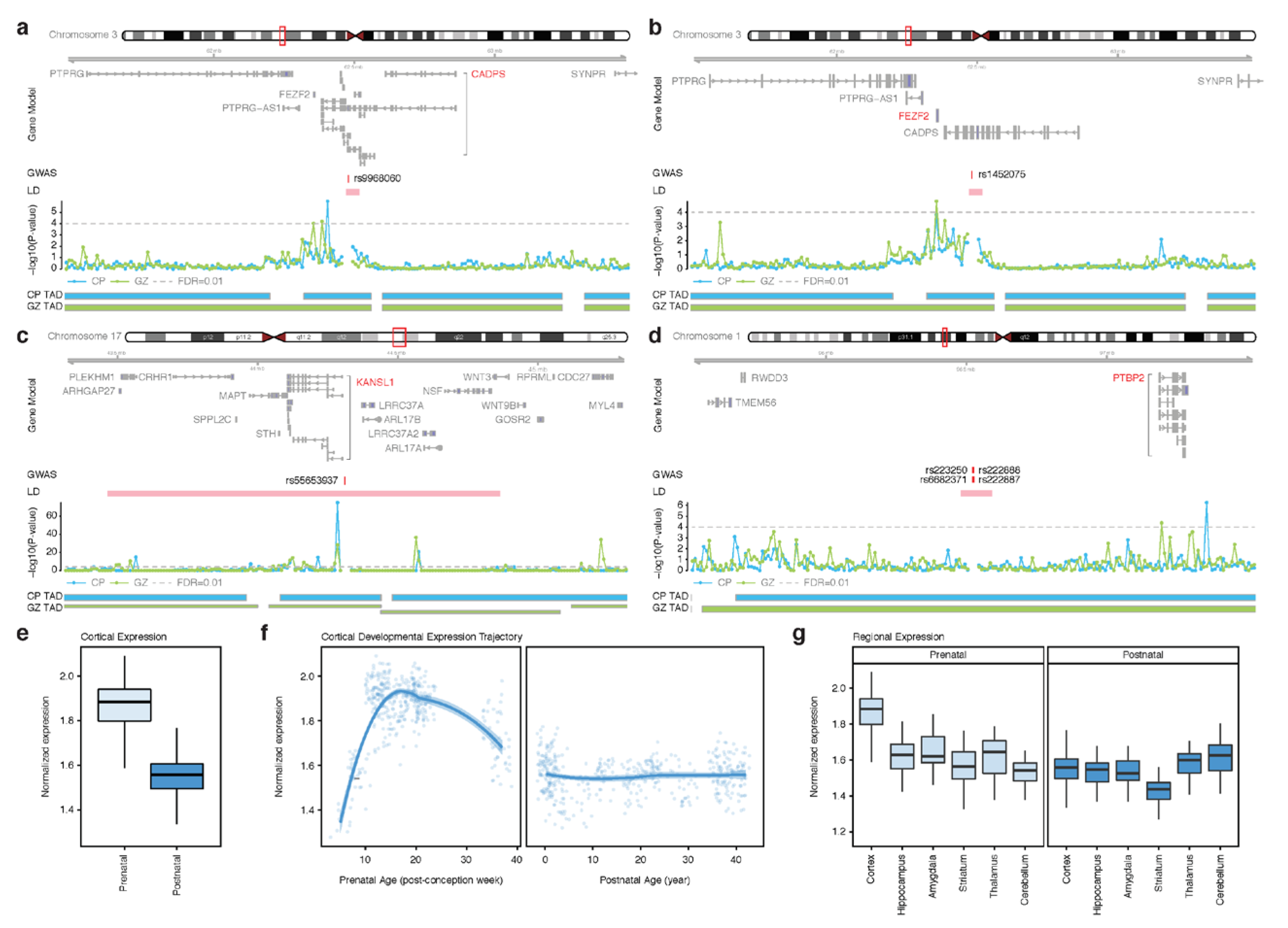
Chromatin interactions identify putative target genes of ASD loci. **a-d**. Chromatin interaction maps of credible SNPs to the 1Mb flanking region, providing putative candidate genes that physically interact with credible SNPs. Gene Model is based on Gencode v19 and putative target genes are marked in red; Genomic coordinate for a credible SNP is labeled as GWAS; – log10(P-value), significance of the interaction between a SNP and each 10kb bin, grey dotted line for FDR=0.01; TAD borders in CP and GZ. **e-f.** Developmental expression trajectories of ASD candidate genes show highest expression in prenatal periods. **g**. ASD candidate genes are highly expressed in the developing cortex as compared to other brain regions.

## Discussion

The high heritability of ASD has been recognized for decades and remains among the highest for any complex disease despite many clinical diagnostic changes over the past 30-40 years resulting in a broader phenotype that characterizes more than 1% of the population. While early GWAS permitted estimates that common polygenic variation should explain a substantial fraction of the heritability of ASD, individually significant loci remained elusive. This was suspected to be due to limited sample size since studies of schizophrenia – with similar prevalence, heritability and reduced fitness – and major depression achieved striking results only when sample sizes five to ten times larger than available in ASD were employed. This study has finally borne out that expectation with definitively demonstrated significant “hits”.

Here we report the first common risk variants robustly associated with ASD by using unique Danish resources in conjunction with results of the earlier PGC data – more than tripling the previous largest discovery sample. Of these, five loci were defined in ASD alone, and seven additional suggested at a stricter threshold utilizing GWAS results from three correlated phenotypes (schizophrenia, depression and educational attainment) and a recently introduced analytic approach, MTAG. Both genome-wide LD score regression analysis and the fact that even among the loci defined in ASD alone there was additional evidence in these other trait scans indicate that the polygenic architecture of ASD is significantly shared with risk to adult psychiatric illness and <u>higher</u> educational attainment and intelligence. It should be noted that the MTAG analyses were carried out as three pairwise analyses. This way we avoid the complex interactions that could arise if we ran three or four correlated phenotypes at a time^9^. Indeed, what we get, despite the secondary summary statistics coming from large, high-powered studies, are relatively modest weights to the contributions from these statistics, because the genetic correlations are modest. The largest weight was 0.27 for schizophrenia, followed by 0.24 for major depression, and 0.11 for educational attainment. Thus all loci identified by MTAG have substantial contributions from ASD alone (**Table 1a, b**) and will most likely also be identified in future ASD-only GWAS with increasing sample sizes.

In most GWAS studies there has been little evidence of heterogeneity of association across phenotypic subgroups. In this study, however, we see strong heterogeneity of genetic overlap with other traits when our ASD samples are broken into distinct subsets. In particular, the excess of alleles associated with higher intelligence and educational attainment was only observed in the higher functioning categories (particularly Asperger’s syndrome and individuals without comorbid ID) – and not in the other/unspecified PDD and ID categories. This is reminiscent, and logically inverted, from the much greater role of spontaneous mutations in these latter categories, particularly in genes known to have an even larger impact in cohorts ascertained for ID/DD^51^. Interestingly, other/unspecified PDD and atypical autism also have a significantly higher PRS for neuroticism than childhood autism and Asperger’s. These different enrichment profiles observed provide evidence for a heterogeneous and qualitatively different genetic architecture between sub-types of ASD, which should inform future studies aiming at identifying etiologies and disease mechanisms in ASD.

The strong differences in estimated SNP heritability between ASD cases with and without ID, and highest in Asperger’s provide genetic evidence of longstanding observations. In particular, this aligns well with the observation that *de novo* variants are more frequently observed in ASD cases with ID compared to cases without comorbid ID, that IQ correlates positively with family history of psychiatric disorders^52^ and that severe ID (encompassing many syndromes that confer high risk to ASD) show far less heritability than is observed for mild ID^53^, intelligence in general^54^ and ASDs. Thus it is perhaps unsurprising that our data suggests that the contribution of common variants may be more prominent in high-functioning ASD cases such as Asperger’s syndrome.

We further explored the functional implications of these results with complementary functional genomics data including Hi-C analyses of fetal brains and brain transcriptome data. Analyses at genome-wide scale (partitioned 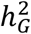 (**Figures S4.3.2-4**) and brain transcriptome enrichment (**Figure 5e-g**)) as well as at single loci (Figure 5a-d, Box 1) highlighted the involvement of processes relating to brain development and neuronal function. Notably, a number of genes located in the identified loci have previously been linked to ASD risk in studies of *de novo* and rare variants (**Box 1**, **Table S3.1.3**), including *PTBP2*, *CADPS*, and *KMT2E*, which were found to interact with credible SNPs in the Hi-C analysis (*PTBP2*, *CADPS*) or contain a loss-of-function credible SNP (*KMT2E*).

Interestingly, aberrant splicing of CADPS’ sister gene CADPS2, which has almost identical function, has been found in autism cases and *Cadps2* knockout mice display behavioral anomalies with translational relevance to autism^55^. *PTBP2* encodes a neuronal splicing factor and alterations in alternative splicing have been identified in brains from individuals diagnosed with ASD^56^.

In summary, we have established a first compelling set of common variant associations in ASD and have begun laying the groundwork through which the biology of ASD and related phenotypes will inevitably be better articulated.

## Methods

### Subjects

#### iPSYCH sample

The iPSYCH ASD sample is a part of a population based case-cohort sample extracted from a baseline cohort^10^ consisting of all children born in Denmark between May 1^st^ 1981 and December 31^st^ 2005. Eligible were singletons born to a known mother and resident in Denmark on their one-year birthday. Cases were identified from the Danish Psychiatric Central Research Register (DPCRR)^12^, which includes data on all individuals treated in Denmark at psychiatric hospitals (from 1969 onwards) as well as at outpatient psychiatric clinics (from 1995 onwards). Cases were diagnosed with ASD in 2013 or earlier by a psychiatrist according to ICD10, including diagnoses of childhood autism (ICD10 code F84.0), atypical autism (F84.1), Asperger’s syndrome (F84.5), other pervasive developmental disorders (F84.8), and pervasive developmental disorder, unspecified (F84.9). As controls we selected a random sample from the set of eligible children excluding those with an ASD diagnosis by 2013.

The samples were linked using the unique personal identification number to the Danish Newborn Screening Biobank (DNSB) at Statens Serum Institute (SSI), where DNA was extracted from Guthrie cards and whole genome amplified in triplicates as described previously^13,57^. Genotyping was performed at the Broad Institute of Harvard and MIT (Cambridge, MA, USA) using the PsychChip array from Illumina (CA, San Diego, USA) according to the instructions of the manufacturer. Genotype calling of markers with minor allele frequency (maf) >0.01 was performed by merging callsets from GenCall^58^ and Birdseed^59^ while less frequent variants were called with zCall^60^. Genotyping and data processing was carried out in 23 waves.

All analyses of the iPSYCH sample and joint analyses with the PGC samples were performed at the secured national GenomeDK high performance-computing cluster in Denmark (https://genome.au.dk).

The study was approved by the Regional Scientific Ethics Committee in Denmark and the Danish Data Protection Agency.

#### Psychiatric Genomic Consortium (PGC) samples

In brief, five cohorts provided genotypes to the sample (n denoting the number of trios for which genotypes were available): The Geschwind Autism Center of Excellence (ACE; N = 391), the Autism Genome Project^61^ (AGP; N = 2272), the Autism Genetic Resource Exchange^62,63^ (AGRE; N = 974), the NIMH Repository (https://www.nimhgenetics.org/available_data/autism/), the Montreal^64^/Boston Collection (MONBOS; N = 1396, and the Simons Simplex Collection ^65,66^(SSC; N = 2231). The trios were analyzed as cases and pseudo controls. A detailed description of the sample is available on the PGC web site: https://www.med.unc.edu/pgc/files/resultfiles/PGCASDEuro_Mar2015.readme.pdf and even more details are provided in Anney et al^5^. Analyses of the PGC genotypes were conducted on LISA on the Dutch HPC center SURFsara (https://userinfo.surfsara.nl/systems/lisa).

#### Follow-up samples

As follow-up for the loci with p-values less than 10^−6^ we asked for look up in 5 samples of Nordic and Eastern European origin with altogether 2,119 cases and 142,379 controls: BUPGEN (Norway: 164 cases and 656 controls), PAGES (Sweden: 926 cases and 3,841 controls not part of the PGC sample above), the Finnish autism case-control study (Finland: 159 cases and 526 controls), deCODE (Iceland 574 cases and 136,968 controls; Eastern Europe: 296 cases and 388 controls). See supplementary for details.

#### GWAS analysis

Ricopili^15^, the pipeline developed by the Psychiatric Genomics Consortium (PGC) Statistical Analysis Group was used for quality control, imputation, principle component analysis (PCA) and primary association analysis. For details see supplementary information. The data was processed separately in the 23 genotyping batches in the case of iPSYCH and separately for each study in the PGC sample. Phasing was achieved using SHAPEIT^16^ and imputation done by IMPUTE2^67,68^ with haplotypes from the 1000 Genomes Project, phase 3^69,70^ (1kGP3) as reference. Chromosome X was imputed using 1000 Genomes Project, phase 1 ^71^ haplotypes.

After excluding regions of high LD^72^, the genotypes were pruned down to a set of roughly 30k markers. See supplementary information for details. Using PLINK’s^73,74^ identity by state analysis pairs of subjects were identified with 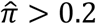 and one subject of each such pair was excluded at random (with a preference for keeping cases). PCA was carried out using smartPCA^75,76^. In iPSYCH a subsample of European ancestry was selected as an ellipsoid in the space of PC1-3 centred and scaled using the mean and 8 standard deviation of the subsample whose parents and grandparents were all known to have been born in Denmark (n=31500). In the PGC sample the CEU subset was chosen using a Euclidian distance measure weighted by the variance explain for each of the first 3 PCs. Individuals more distant than 10 standard deviations from the combined CEU and TSI HapMap reference populations were excluded. We conducted a secondary PCA to provide covariates for the association analyses. Numbers of subjects in the data generation flow for the iPSYCH sample are found in the descriptive table **Table S1.1.1**.

Association analyses were done by applying PLINK 1.9 to the imputed dosage data (the sum of imputation probabilities P(A1A2) + 2P(A1A1)). In iPSYCH we included the first four principal components (PCs) as covariates as well as any PC beyond that, which were significantly associated with ASD in the sample, while the case-pseudo-controls from the PGC trios required no PC covariates. Combined results for iPSYCH and for iPSYCH with the PGC was achieved by meta-analysing batch-wise and study-wise results using METAL^77^ (July 2010 version) employing an inverse variance weighted fixed effect model^22^. On chromosome X males and females were analyzed separately and then meta-analyzed together. Subsequently we applied a quality filter allowing only markers with an imputation info score ≥ 0.7, maf ≥ 0.01 and an effective sample size (see supplementary info) of at least 70% of the study maximum. The degree to which the deviation in the test statistics can be ascribed to cryptic relatedness and population stratification rather than to polygenicity was measured from the intercept in LD score regression^23^ (LDSC) as the ratio of (intercept-1) and (mean(χ^2^)-1).

MTAG^9^ was applied with standard settings. The iPSYCH-PGC meta-analysis summary statistics was paired up with the summary statistics for each of major depression^29^ (excluding the Danish sampled but including summary statistics from 23andMe^78^, 111,902 cases, 312,113 controls, and mean χ^2^=1.477), schizophrenia^15^ (also excluding the Danish samples, 34,129 cases, 45,512 controls, and mean χ^2^=1.804) and educational attainment^26^ (328,917 samples and mean χ^2^=1.648). These are studies that have considerably more statistical power than the ASD scan, but the genetic correlations are modest in the context of MTAG, so the weights ascribed to the secondary phenotypes in the MTAG analyses remain relatively low (no higher than 0.27). See supplementary methods for details.

The results were clumped and we highlighted loci of interest by selecting those that were significant at 5 x 10^−8^ in the iPSYCH-PGC meta-analysis or the meta-analysis with the follow-up sample or were significant at 1.67 x l0^-8^ in any of the three MTAG analyses. The composite GWAS consisting of the minimal p-values at each marker over these five analyses wsa used as a background when creating Manhattan plots for the different analyses showing both what is maximally achieved and what the individual analysis contributes to that.

#### Gene-based association and gene-set analyses

MAGMA 1.06 ^32^ was applied to the summary statistics to test for gene-based association. Using NCBI 37.3 gene definitions and restricting the analysis to SNPs located within the transcribed region, mean SNP association was tested with the sum of -log(SNP p-value) as test statistic. The resulting gene-based p-values were further used in competitive gene-set enrichment analyses in MAGMA. One analysis explored the candidate sets M13, M16 and M17 from Parikshak et al. 2013 ^33^ as well as constrained, loss-of-function intolerant genes (pLI>0.9)^34,35^ derived from data of the Exome Aggregation Consortium (see supplementary methods for details). Another was an agnostic analysis of the Gene Ontology sets^37,38^ for molecular function from MsigDB 6.0 ^39^. We analyzed only genes outside the broad MHC region (hg19:chr6:25-35M) and included only gene sets with 10-1000 genes.

#### SNP heritability

SNP heritability, 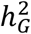, was estimated using LDSC^23^ for the full sample and GCTA^40–42^ for subsamples too small for LDSC. For LDSC we used precomputed LD scores based on the European ancestry samples of the 1000 Genomes Project^71^ restricted to HapMap3^79^ SNPs (downloaded from the https://github.com/bulik/ldsc). The summary stats with standard LDSC filtering were regressed onto these scores. For liability scale estimates, we used a population prevalence for Denmark of 1.22%^21^. Lacking proper prevalence estimates for subtypes, we scaled the full spectrum prevalence based on the composition of the case sample.

For subsamples too small for LDSC, the GREML approach of GCTA^40–42^ was used. On best guess genotypes (genotype probability >0.8, missing rate < 0.01 and MAF > 0.05) with INDELs removed, a genetic relatedness matrix (GRM) was fitted for the association sample (i.e. the subjects of European ancestry with 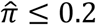) providing a relatedness estimate for all pairwise combinations of individuals. Estimation of the phenotypic variance explained by the SNPs (REML) was performed including PC 1-4 as continuous covariates together with any other PC that was nominally significantly associated to the phenotype as well as batches as categorical indicator covariates. Testing equal heritability for non-overlapping groups was done by permutation test (with 1000 permutations) keeping the controls and randomly the assigning the different case labels.

Following Finucane et al^47^, we conducted an enrichment analysis of the heritability for SNPs for functional annotation and for SNPs located in cell-type-specific regulatory elements. Using first the same 24 overlapping functional annotations (stripped down from 53) as in Finucane et al. we regressed the *χ*^2^ from the summary statistics on to the cell-type specific LD scores download from the site mentioned above with baseline scores, regression weights and allele frequencies based on European ancestry 1000 Genome Project data. The enrichment of a category was defined as the proportion of SNP heritability in the category divided by the proportion of SNPs in that category. Still following Finucane et al. we did a similar analysis using 220 cell type–specific annotations divided into 10 overlapping groups. In addition to this, we conducted an analysis based on annotation derived from data on H3K4Me1 imputed gapped peaks data from the Roadmap Epigenomics Mapping Consortium^80,81^; more specifically information excluding the broad MHC-region (chr6:25-35MB).

#### Genetic correlation

For the main samples, SNP correlations, *r_G_*, were estimated using LDSC^23^ and for the analysis of ASD subtypes and subgroups where the sample size were generally small, we used GCTA^42^. In both cases, we followed the same procedures as explained above. For all but a few phenotypes, LDSC estimates of correlation were achieved by uploaded to LD hub^25^ for comparison to all together 234 phenotypes.

#### Polygenic risk scores

For the polygenic risk scores (PRS) we clumped the summary stats applying standard Ricopili parameters (see supplementary methods for details). To avoid potential strand conflicts we excluded all ambiguous markers for summary statistics not generated by Ricopili using the same imputation reference. PRS were generated at the default p-value thresholds (5e-8, 1e-6, 1e-4, 0.001, 0.01, 0.05, 0.1, 0.2, 0.5 and 1) as a weighted sum of the allele dosages. Summing over the markers abiding by the p-value threshold in the training set and weighing by the additive scale effect measure of the marker (log(OR) or β) as estimated in the training set. Scores were normalized prior to analysis.

We evaluated the predictive power using Nagelkerke’s *R*^2^ and plots of odds ratios and confidence intervals over score deciles. Both *R*^2^ and odds ratios were estimated in regression analyses including the relevant PCs and indicator variable for genotyping waves.

Lacking a large ASD sample outside of iPSYCH and PGC, we trained a set of PRS for ASD internally in the following way. We divided the sample in five subsamples of roughly equal size respecting the division into batches. We then ran five GWAS leaving out each group in turn from the training set and meta-analyzed these with the PGC results. This produced a set of PRS for each of the five subsamples trained on their complement. Prior to analyses, each score was normalized on the group where it was defined. We evaluated the predictive power in each group and on the whole sample combined.

To exploit the genetic overlap with other phenotypes to improve prediction, we created a series of new PRS by adding to the internally trained ASD score the PRS of other highly correlated phenotypes in a weighted sum. See supplementary info for details.

To analyze ASD subtypes in relation PRS we defined a hierarchical set of phenotypes in the following way: First hierarchical subtypes was childhood autism, hierarchical atypical autism was defined as everybody with atypical autism and no childhood autism diagnosis, hierarchical Asperger’s as everybody with an Asperger’s diagnosis and neither childhood autism nor atypical autism. Finally we lumped other pervasive developmental disorders and pervasive developmental disorder, unspecified into pervasive disorders developmental mixed, and the hierarchical version of that consists of everybody with such a diagnosis and none of neither preceding ones (**Table S2.3.1**). We examined the distribution over the distinct ASD subtypes of PRS for a number of phenotypes showing high *r_G_* with ASD (as well as a few with low *r_G_* as negative controls), by doing multivariate regression of the scores on the subtypes while adjusting for relevant PCs and wave indicator variables in a linear regression. See supplementary methods for details.

#### Hi-C analysis

The Hi-C data was generated from two major cortical laminae: the germinal zone (GZ), containing primarily mitotically active neural progenitors, and the cortical and subcortical plate (CP), consisting primarily of post-mitotic neurons^49^. We first derived a set of credible SNPs (putative causal SNPs) from the identified top ranking 29 loci using CAVIAR^50^. To test whether credible SNPs are enriched in active marks in the fetal brain^81^, we employed GREAT as previously described^49,82^. Credible SNPs were, sub-grouped into those without known function (unannotated) and functionally annotated SNPs (SNPs in the gene promoters and SNPs that cause nonsynonymous variants) (**Figure S4.5.1**). Then we integrated unannotated credible SNPs with chromatin contact profiles during fetal corticogenesis^49^, defining genes physically interacting with intergenic or intronic SNPs (**Figure S4.5.1**).

Spatiotemporal transcriptomic atlas of human brain was obtained from Kang et al^83^. We used transcriptomic profiles of multiple brain regions with developmental epochs that span prenatal (6-37 post-conception week, PCW) and postnatal (4 months-42 years) periods. Expression values were log-transformed and centered to the mean expression level for each sample using a *scale(center=T, scale=F)+1* function in R. ASD candidate genes identified by Hi-C analyses (Figure S4.5.1) were selected for each sample and their average centered expression values were calculated and plotted.

## Summary statistics

The summary statistics are available for download the iPSYCH download page http://ipsych.au.dk/downloads/ and at the PGC download site: https://www.med.unc.edu/pgc/results-and-downloads.

## Acknowledgements

The iPSYCH project is funded by the Lundbeck Foundation (grant numbers R102-A9118 and R155-2014-1724) and the universities and university hospitals of Aarhus and Copenhagen. Genotyping of iPSYCH and PGC samples was supported by grants from the Lundbeck Foundation, the Stanley Foundation, the Simons Foundation (SFARI 311789 to MJD), and NIMH (5U01MH094432-02 to MJD). The Danish National Biobank resource was supported by the Novo Nordisk Foundation. Data handling and analysis on the GenomeDK HPC facility was supported by NIMH (1U01MH109514-01 to Michael O’Donovan and ADB). High-performance computer capacity for handling and statistical analysis of iPSYCH data on the GenomeDK HPC facility was provided by the Centre for Integrative Sequencing, iSEQ, Aarhus University, Denmark (grant to ADB). Dr SD Rubeis was supported by NIH grant MH097849 (to JDB), MH111661 (to JDB), and the Seaver Foundation (to SDR and JDB). Dr J Martin was supported by the Wellcome Trust (grant no: 106047). We thank the research participants and employees of 23andMe for making this work possible.

## Members of the PGC-MDD Group and the 23andMe Research Team

See list in the supplementary file.

## Author contributions

Analysis: JG, SR, TDA, MM, RW, HW, JP, FB, JHC, CC, KD, SDR, BD, SD, MEH, SH, DH, HH, LK, JM, JM, ARM, MN, TN, DSP, TP, BSP, PQ, JR, EBR, KR, PR, ES, SS, FKS, SS, PFS, PT, GBW, XX, DG.

JG, BMN, MJD, and ADB supervised and coordinated the analyses.

Sample and/or data provider and processing: JG, SR, MM, RW, EA, OA, RA, RB, JDB, JBG, MBH, FC, KC, DD, AD, JG, CH, MVH, CH, JM, AP, CP, MGP, JBP, KR, AR, GDS, HS, KS, CS, PGC-ASD, BUPGEN, PGC-MDD, 23andMeResearch Team, DH, OM, PBM, BMN, MJD, ADB.

Core PI group: DG, MN, DH, TW, OM, PBM, BMN, MJD, ADB.

Core writing group: JG, MJD, ADB.

Direction of study: MJD, ADB.

